# We think what we eat: Animal-based diet influences cerebral and microbiota networks connectivity in early ages. A study case of an indigenous community in Mexico

**DOI:** 10.1101/2020.07.25.221408

**Authors:** Ramirez-Carrillo Elvia, G-Santoyo Isaac, López-Corona Oliver, Olga A. Rojas-Ramos, Luisa I. Falcón, Osiris Gaona, Daniel Cerqueda-García, Andrés Sánchez-Quinto, Rosa María de la Fuente Rodríguez, Ariatna Hernández Castillo, Nieto Javier

## Abstract

We are not individuals, we are much better described as ecosystems due to trillions of bacteria and other microorganisms that inhabit us. We now know that gut microbiota can greatly influence many physiological parameters that in turn may impact several cognitive functions, such as learning, memory, and decision making processes. This mutualistic symbiotic relation known as the gut-brain axis is also constrained by external factors such as dietary habits such as animal protein and lipids intake. Using a novel combination of Machine Learning and Network Theory techniques, we provide evidence from an indigenous population in Guerrero Mexico, that both brain and gut-microbiota connectivity, evaluated by Minimum Spanning Tree as the critical backbone of information flow, diminish under either low protein or lipids intake. We discuss then how this loss of connectivity may translate into a reduction of the individual’s capacity to cope with perturbations as loss of connectivity may be linked with losses in antifragility.

## Introduction

The gut-brain axis is an integrative physiological mechanism that connects endocrine, immunological, dietary, and neuronal signals between the gut system and the brain (Mayer 2011). In that sense, we know that diet plays a crucial role in both the ecosystem dynamics of gut microbiota (GM) and brain functioning. Hence, even temporary changes in diet habits may cause important effects on the gut-brain axis functioning (Sandhu et al 2017). In this work, we show how the consumption of animal proteins and lipids impacts the bacteria GM and neural connectivity, using a combination of field and laboratory work with novel network theory and Machine Learning methods.

It is well documented that the incorporation of animal products such as protein and lipids in the hominid diet was one of the major evolutionary processes driving human evolution. Some three million years ago, another time plentiful food supply, savannahs went through a severe climate change that dried them up. Under those severe new conditions, two types of primates survived the catastrophe, the ones with a plant-processing machine and the ones with a meat-eating machine. The second ones became us by evolving a bigger and more complex brain enhanced by the new supply of iron, zinc, vitamin B12, and fatty acids among other key nutrients (Gupta, 2016). Although some of these nutrients are also present in plants, animal meat is rich for example in hemoglobin-derived iron, which is absorbed more easily than the non-haem type found in beans and leafy greens. In addition, compounds known as phytates in plants bind to the iron and obstruct its availability to the body. Consequently, meat is a much richer dietary iron source than any plant food.

One of the evolutionary consequences of this dietary transition could have resulted in what Aiello and Wheeler (1995) named as the expensive tissue hypothesis, which essentially states that since both intestine and brain are composed of metabolically expensive tissues, which need a disproportionate amount of energy to function properly, then a large brain would be incompatible with a large intestine, associated with a plant-based diet. This simple trade-off translates into smaller guts with the same or more energy needs (as the brain gets larger), so the only way to make it work is by modifying the diet to one that offers high nutritional rates that is easy to handle, that is meat (Pobiner, 2016).

In this way, our gut, brain, and diet have co-evolved with the animals we eat and the technology we used (i.e. fire and cooking) in what we have called the ecobiont ontology (López-Corona et. al. 2019). Even more, this implies that our GM also has co-evolved in a meat-based diet and so one would expect that animal protein consumption may have an observable effect on the gut-brain axis structure and functioning.

In this line of thought, evidence from murine models and human studies have demonstrated that animal protein consumption modulates important events in the development of the central nervous system such as neuronal proliferation, axon and dendrite growth, synaptogenesis and neuronal apoptosis (Allamy & Bengelloun, 2012; Prado & Dewey, 2014). As happens with protein intake; myelination, another essential and required event in neurodevelopment is influenced by a second dietary animal product, lipids. For instance, brain lipids specify the position and role of proteins in the cell membrane and thus control synaptic neuron signals (Shamim et. al. 2018). Lipids may also act as transmitters and relay signals to intracellular compartments or other cells from the membrane (Shamim et al., 2018).

Just like the brain, GM is another system that is profoundly influenced by the host’s dietary patterns (De Filippo et. al. 2010). This is important because the emergence of Next-Generation DNA sequencing methods has led to a renewed appreciation of the importance of GM as a regulatory component of brain functioning and vice-versa (Cryan et. al. 2019). Nevertheless, while the composition and abundance of our GM are sensible to dietary changes, determined for example by substratum competition and intestinal tolerance; predicting the impact of diet on a complex and interactive system like the human GM needs not only more knowledge about the component groups involved but also more integration of such knowledge (Flint et. al. 2015). Interestingly, the maturation of both systems occurs in parallel and they have similar critical developmental windows, being childhood and adolescence the most dynamic period for this communication, and where any disruption has the potential to profoundly alter brain-gut signaling (Borre et. al., 2014).

This points in the direction of a complex systems perspective, in which it is not only important to characterize the elements that constitute the systems, but above all its interactions. For example, in recent work, it has been suggested that behavioral changes, such as the development and persistence of depression, may result from a complex network of communication between macro-and micro-organisms capable of modifying different host’s neurophysiological components (Ramírez-Cariillo et. al. 2020). So it should be self-evident that if we pretend to understand brain connectivity, we need to talk about GM connectivity too.

In terms of a complex systems approach, in a recent paper by Saba and coworkers (2019) it has been summarized how even when Network Theory methods (one of the main streams in complexity) are becoming widespread (Bullmore and Sporns, 2009; Drakesmith et al., 2015; Chiang et al., 2016; Agosta et al., 2013; Filippi et al., 2017) and have shown to be useful in the study of for example of brain dysfunction mechanism (Bullmore and Sporns, 2012); a number of subjective choices significantly hamper the methodology. Furthermore, several network metrics, including node centrality indices assume different significance on a local or global scale (Telesford et al., 2011; Antonenko et al., 2018). In that respect the authors make the case for the use of Minimum Spanning Tree as an unambiguous solution, ensuring accuracy, robustness, and reproducibility that also avoids methodological confounding thanks to the way it takes both topological properties and functional connectivity information into account (Tewarie et al., 2014; van Diessen et al., 2016; van Lutterveld et al., 2017).

The minimum spanning tree or MST problem is one of the best-studied optimization problems in computer science. The task is to calculate a tree that connects all the vertices of a network (a spanning tree) using only edges in such a way that total weight is minimum among all possible spanning trees (Eisner 1997). But perhaps the most interesting aspect of MST measurements is that if we consider the brain network as an information transport network in which cerebral areas are considered nodes, and their connections as edges (Rubinov and Sporns, 2010), it enables both structural (anatomical) and functional (statistical relationship between two nodes) connectivity to be assessed (Zhang et al., 2016). Then an MST might represent the critical backbone of information flow (Saba et.al. 2019).

This kind of Network Theory analysis has been getting a lot of attention recently because it opens a multidisciplinary approach that allows us to analyze the brain as a complex system in a straightforward computable way (Rubinov and Sporns, 2010) and not merely as a metaphor. In the same way, we have pointed out in previous work about the disturbance in human gut microbiota networks by parasites (Ramírez-Carrillo et. al. 2020), that the novelty of these analyses is that they allow us to study the interactions allowing us to see the whole system but also how different components are affected.

In this work, we will follow this Network Theory approach to study whether the consumption of animal protein and lipids in children can influence the connectivity of both, brain neocortex activity measured by quantitative electroencephalography (EGG), and GM determined by 16S rDNA gene high-throughput sequencing. To accomplish this, we have selected an indigenous Mexican community belonging to the ethnic group Me’phaa in the state of Guerrero. In this community, the consumption of these animal-based nutrients is highly limited (Borda-Niño et al., 2016). Hence, we hypothesized that inter-individual variations in the access of these limiting nutrients would have a greater effect on the connectivity of both axes. Moreover, this community also show close inter-individual similarities in relation to important factors that influence Brain activity and GM, such as the use of allopathic medication, access to health services, economic status, parent and child scholarly and access to computational and communications technologies (Sonnenburg and Sonnenburg, 2019; Lipina and Evers, 2017). Hence, this human population offers us a unique model that controls these factors in a natural and ecologically valid way.

## Methods

### Study subjects

The population selected in this study corresponds to a pre-hispanich indigenous community from the etnich group Me Phaa located in a region named “La Montaña Alta” in the state of Guerrero, México. It is an interesting case of study because, on the one hand, it is one of the more contrasting groups in terms of lifestyle concerning typical westernized urbanized areas (Black, 2014; Camacho, 2007; Miramontes et.al. 2012). On the other hand, because isolation, tradition and poverty conditions, their diet is provided basically by subsistence farming; recollection of wild edible plants; consumption of some fruits and vegetables cultivated in garden plots; and meanwhile animal-based products are available by hunting, it is consumed almost entirely during special occasions and is not part of the daily diet (Borda-Niño et.al. 2016). So their intake of protein and lipids is below the national mean and is very homogenous inside the population. The present study is composed by a sample size of 33 indigenous children between 5-10 years old (12 males and 19 females; Table 1).

**Table 1.**
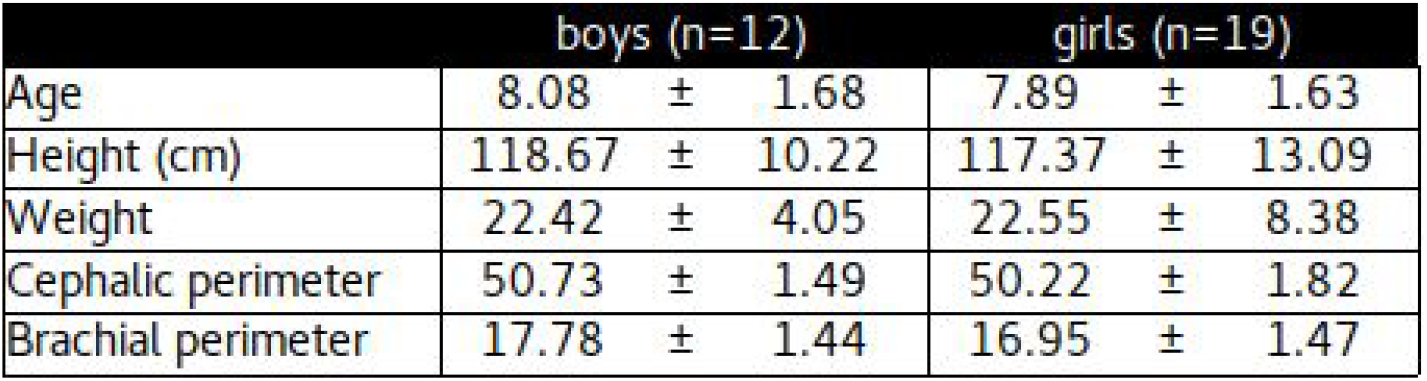
Anthropometric Measurements of study samples

### Anthropometric Measurements

Standardized basic anthropometric measurement protocols were followed. These included weight measurement for each participant using an OMRON brand digital scale (HBF-514C), as well as their height using a portable leveled stadiometer (HM200P, PortStad) placed on the wall at 2m height. Also, the head and brachial circumference were taken with a measuring tape (Seca ™).

### Gut microbiota composition

#### Sample collection

GM bacteria composition was obtained from fecal samples using a 16S rDNA High-throughput sequencing. To accomplish this, each fecal sample was collected in a sterilized plastic jar with an unique identifier written on the lid. After recollection, all samples were frozen with liquid nitrogen to a posterior storage at −20°C until DNA extraction.

#### Fecal DNA extraction

Each DNA sample (~100 μl of fecal DNA) was extracted using the DNeasy Blood & Tissue kit (Qiagen, Valencia, CA) according to the manufacturer’s instructions. DNA was resuspended within 30 μl of molecular grade water and stored at −20°C until PCR amplification.

#### 16S rDNA gene amplification and sequencing

For the 16S rDNA gene amplification and sequencing we used the hypervariable V4 region of the 16S rDNA gene with universal bacterial/archaeal primers 515F/806R following the procedures reported by Caporaso et al. (2012) 70. The PCR mix was carried out in 25 μl reactions by triplicate per samples as follows: 2.5 μl Takara ExTaq PCR buffer 10X, 2 μl Takara dNTP mix (2.5 mM), 0.7 μl bovine serum albumin (BSA, 20 mg ml-1), 1 μl primers (10 μM), 0.125 μl Takara Ex Taq DNA Polymerase (5 U μl-1) (TaKaRa, Shiga, Japan), 2 μl of DNA and 15.67 μl nuclease-free water. The PCR protocol included an initial denaturation step at 95°C (3 min), followed by 35 cycles of 95°C of 30 s, 52°C of 40 s, and 72° C of 90 s; followed by a final extension of 12 min at 72°C. Triplicates were pooled and purified using the SPRI magnetic bead, AgencourtAMPure XP 214 PCR purification system (Beckman Coulter, Brea, CA, USA). The characterization of the fecal purified 16S rDNA fragments (~20 ng per sample) were sequenced on an IlluminaMiSeq platform (Yale Center for Genome Analysis, CT, USA), generating ~250 bp paired-end reads. All sequences obtained were uploaded to the NCBI database under the Bioproject number PRJNA593240.

#### Analysis of the sequence data

The paired-end 2×250 reads were processed in QIIME2. The reads were denoised with the DADA2 plugin to resolve the amplicon sequence variants (ASVs) (Callahan et.al. 2016). Both forward- and reverse-reads were truncated at 200 pb, and chimeric sequences were removed using the “consensus” method. Representative ASVs sequences were taxonomically assigned using the “classify consensus-vsearch pluggin” (Rognes et.al. 2016), using the SILVA 132 database as a reference (Quast et.al. 2013). An alignment was performed with the MAFFT algorithm (Katoh et.al. 2009). After masking positional conservations and gap filtering, a phylogeny was built with the FastTree algorithm (Price et.al. 2010). The abundance table and phylogeny were exported to the R environment to perform the statistical analysis with the phyloseq (McMurdie et.al. 2013) and ggplot2 packages. Plastidic ASVs were filtered out of the samples, which were rarefied to a minimum sequencing effort of 21 000 reads per sample. The total diversity (alpha diversity) of the ASVs was calculated using Faith’s Phylogenetic Diversity Index (PD), Shannon’s Diversity Index and Observed ASV’s.

### Proteins and Lipids intake estimation

To evaluate the amount of proteins and lipids that children consume, we applied the food frequency questionnaire (FFQ) from the mexican national health system ENSANUT-2012 (https://ensanut.insp.mx/), which was adapted for available food in communities on the study site. The ENSANUT uses a probabilistic multistage stratified cluster sampling design to be representative of the country and regions. The FFQ was answered for the mother of every child registering the composition of the child’s diet by counting the number of times any given food was consumed per day and per week, obtaining monthly food intake. Once consumption frequency was registered, an approximate monthly intake of animal fat and protein grams were calculated according to portions and based on the Mexican System of Equivalent Foods (SMAE, 2014). For protein and lipids, the consumption of milk, yogurt, pork, beef, chicken, egg, butter, and fish was taken into account in order to standardize the record of consumption of these foods and their comparison with international standards. Approximate daily protein and lipid consumption was calculated by dividing the monthly consumption by 30 days.

### Brain Cortex Activity

In order to obtain an estimate of the child’s brain cortex activity we obtained the absolute power during a resting-state period using quantitative EEG. Standardized preparation procedures were followed using GRAEL 4K EEG Amplifier, electrode caps and the acquisition program Profusion EEG5. For the acquisition process, small (50-54 cm) and medium (54-58 cm) size Electrocap universal caps at 19 localitations according to the 10-20 International System (Jasper,1958) were used. The caps were chosen depending on cephalic circumference. A resting-state EEG record through a monopolar montage using as reference each ipsilateral ear lobe was acquired, and an extra electrode at the external canthus of the left eye to record eye movements was placed.

### Signal acquisition and processing

Data was recorded with a bandwidth of 1 Hz to 70 Hz, an on-line Notch filter and a sampling rate of 512 Hz; impedances below 10 kΩ were maintained in all the electrodes; a 5-minute open-eye and 5-minute closed-eye resting state recording was performed for each participant. The recording was done using a counterbalance between individuals to start with eyes open or eyes closed depending on the subjects’ disposition. In each procedure, 15-20 minutes were recorded, switching conditions to achieve 2 uninterrupted minutes of each condition (eyes open and eyes closed).

An offline visual inspection was performed in the Profusion EEG5 to obtain 2 minutes running without artifacts of each condition. Data was analyzed using a personalized script in Matlab and the Fieldtrip toolbox, filters above 1 Hz and below 40 Hz was applied, a variance-based distribution method between trials and a second visual inspection was performed to obtain 39 to 42 second artifact-free epochs of each subject and each condition. Finally, the data was transformed to logarithmic scale to proceed with the power analysis. The absolute power was obtained by Fourier Analysis for each derivation, participant and condition, by classic broad bands Delta (1-3 Hz), Theta (4-7 Hz), Alpha (8-14 Hz) and Beta (15-30 Hz). Three electrodes per brain region of interest (ROI’s) as means of testing hypotheses of brain function were averaged to create five nodes. ROIS’s contained the following electrodres; left anterior (Fp1, F3, F7), right anterior (Fp2, F4, F8), left posterior (P3,T7, O1), right posterior (P4, T8, O2) and midline (Fz, Cz, Pz).

### Network analysis for EEG and GM data

For all analyses considering the two variables of interest: Protein and lipids intake; we divided the population (the 33 children) into two categories, Low or High, if the individual intake was below or above the mean intake in the population. In order to have a balanced number of individuals inside each category for protein intake, we use as threshold z values of +/- 0.5 for High and Low intake.

Functional connectivity in the human brain can be represented as a network using electroencephalography (EEG) data (Rathee et al., 2017). It was common practice to use correlation measurements to construct those networks, but both theory and practice shows that Mutual Information (MI) is a better way of doing it. On the one hand we know that entropy-based metrics, such as MI are vastly more potent than correlation and particularly that MI can detect nonlinearities, it is additive and ergodic (Taleb, 2020). On the other hand, Zhang and co-workers (2018) strongly conclude from their study of children with epilepsy that functional connectivity measured using MI captures physiologically relevant features of the brain network better than correlation. In the same way, there are several applications of connectivity measured by MI networks such as Sun and co-workers (2018) with in post-stroke patients with different levels of depression, or as a useful biomarker for preclinical Alzhaimer (Gaubert et.al., 2019).

Mutual Information calculations were performed using the ‘mpmi’ R Package (https://cran.r-project.org/web/packages/mpmi/mpmi.pdf) which uses a kernel smoothing approach to calculate Mutual Information for comparisons between all types of variables including continuous vs continuous, continuous vs discrete and discrete vs discrete. The final result from mpmi is a MI matrix equivalent to a standard correlation.

Once MI matrix is obtained from EEG data, we construct the corresponding networks using MI coefficients as weights in Cytoscape (Shannon et. al. 2003), considering for these five nodes: front left, front right; posterior left, posterior right; and midline.

Afterward, we calculate the Minimum Spanning Tree (MST) in Gephi (Bastian et. al. 2009) which finds an edge of the least possible weight that connects any two trees in the forest. This algorithm finds a subset of the edges that forms a tree that includes every vertex, where the total weight of all the edges in the tree is minimized. In our context as each weight is a MI-connectivity value, the weight of the MST (the sum of the MST’s weights) is a measure of both structural and functional brain connectivity (Tewarie et.al., 2015).

As not every species participate informationally into protein or lipid intake differences, we used the Machine Learning algorithm of Random Forest (Breiman, 2001) implemented in R under the randomForest package (https://cran.r-project.org/web/packages/randomForest/randomForest.pdf), to identify main bacteria families (nodes) in terms of its Mean Decrease Gini value which tell us the relative importance of the node for the classification process in terms of High Vs Low intake. With these families identified, using the dataset of ASVs (i.e. bacteria species) relative abundances in fecal samples, we used the Cooccur package (https://cran.r-project.org/web/packages/cooccur/cooccur.pdf) in R74 to construct a co-occurrence matrix and use it as weight in microbiota network.

Finally we construct the corresponding subnetworks and calculate the MST values as done for the EEG data.

## Results

Figure 1 shows the EEG mutual information matrices. Here, lines represent the level of mutual information in the corresponding networks of each broad brain band and two conditions, open and closed eyes.

**Figure 1.**
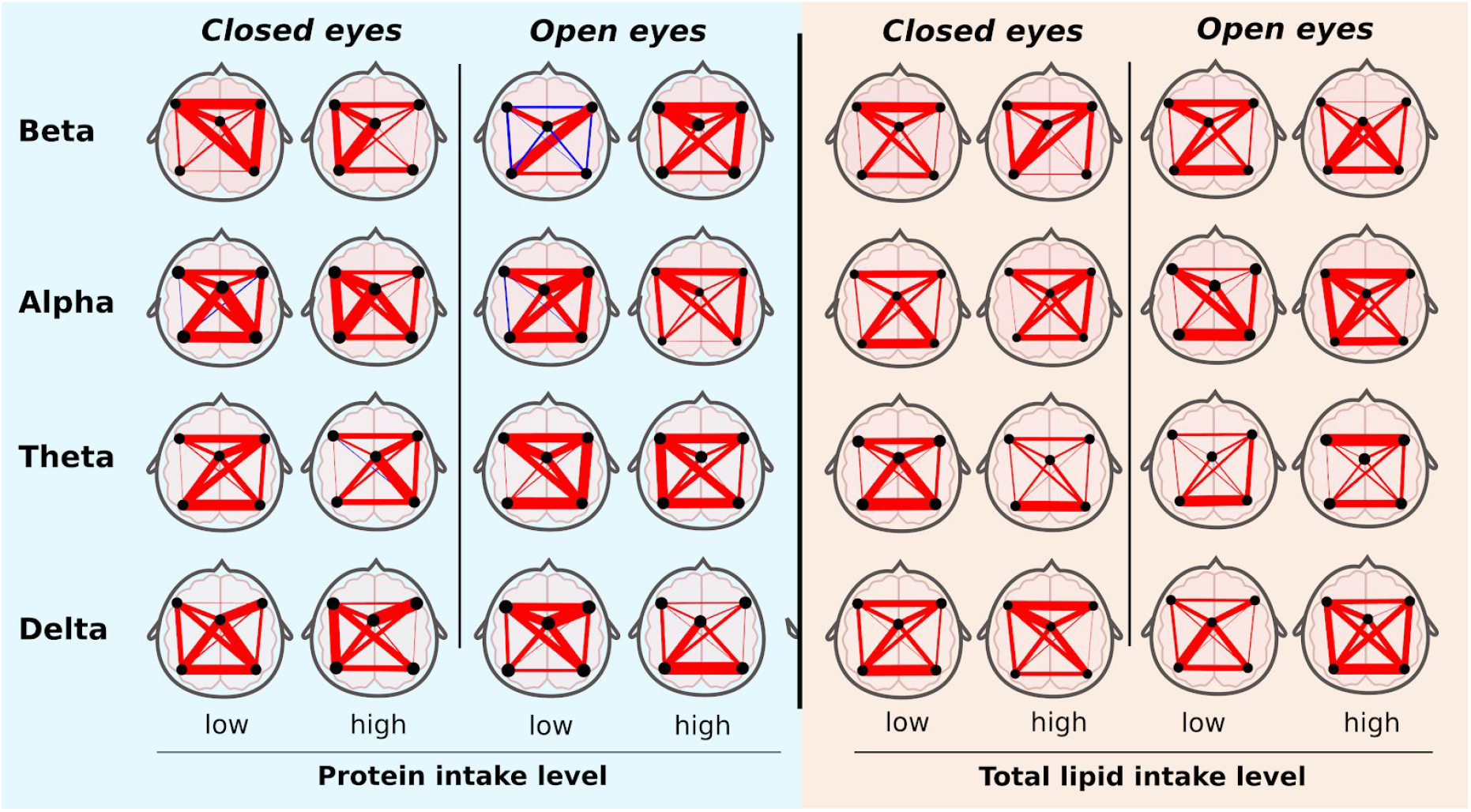
We show a resting-state EEG Mutual Information networks, considering five nodes: front left, front right; posterior left, posterior right; and midline. Mutual Information was obtained for each broad brain band and resting-state condition; Open and Closed eyes. The color red means positive relationship, blue negative, just as in a standard correlation. The thickness of the line represents the intensity of the relationship.

In general, there are few negative relationships (i.e. Blue lines in Fig. 1). Qualitatively, there are evident differences between networks of distinct bands and between levels of protein intake. In particular, it is interesting to see qualitatively the contrast in open eyes when the beta band is evaluated. Here, a low protein intake results in negative mutual information between left, right, anterior and posterior nodes, while with high intake there was a positive and very strong mutual information. In addition, the high intake of lipids would seem to connect the posterior and middle regions of the brain.

In order to make a more quantitative discussion of the effect of diet on brain connectivity, the MST was calculated following what was done by Saba and co-workers (2019). The results of this analysis can be seen in Figure 2 where with the exception for the beta band, it is observed that in the open eye state, which represents the condition of the highest cognitive demand, there is a systematic increase in brain connectivity for all bands in the highest protein and lipid groups.

**Figure 2.**
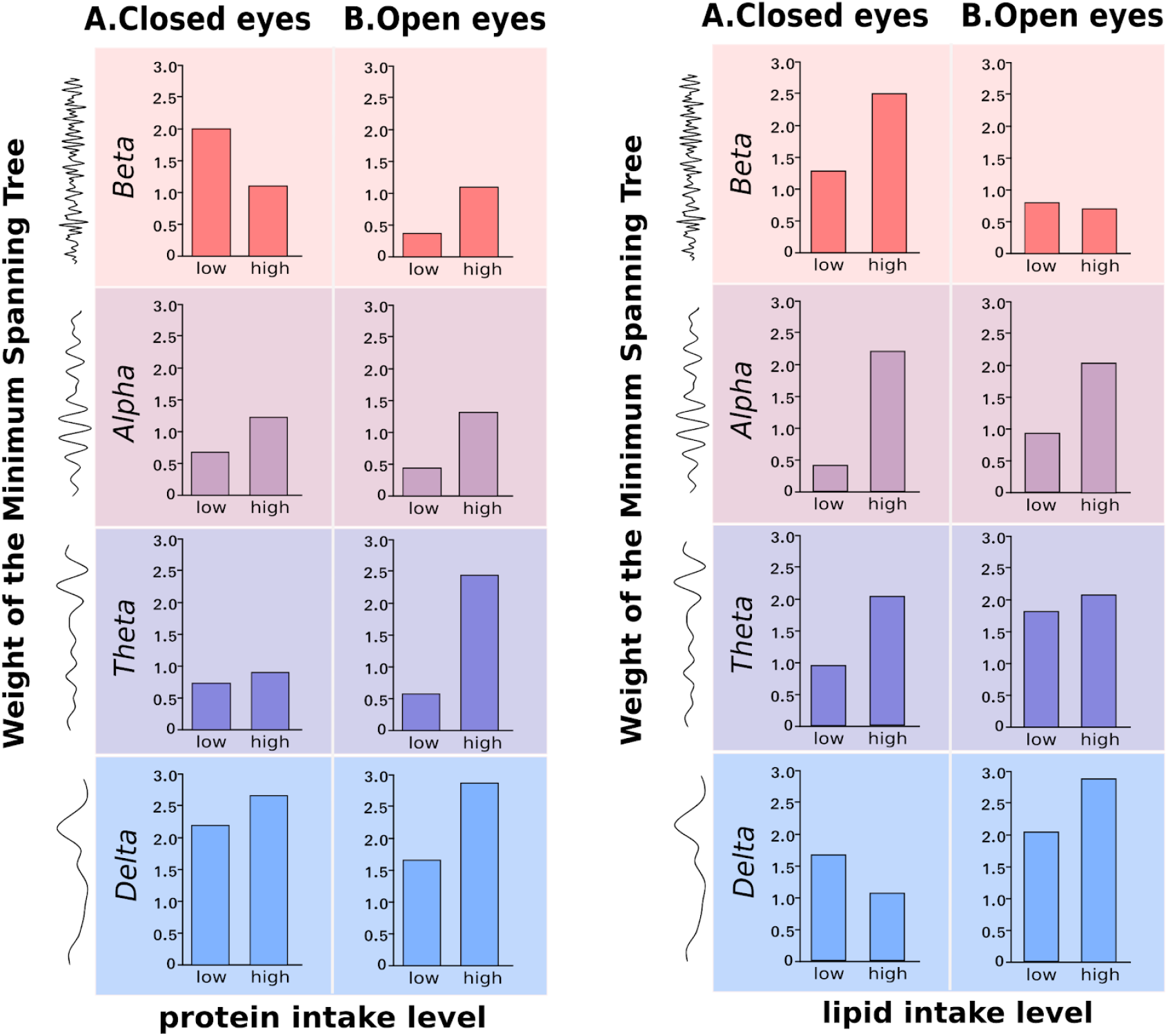
Minimum spáning tree (MST) of absolute brain power networks for protein or lipid intake level in both condition A) closed or B) Open eyes. Weight of the MST as a measure of total structural and functional network connectivity (the higher the value, the higher connectivity).

To assess GM connectivity through Minimum Spanning Tree (MST), we first performed a standard classification analysis by calculating a Random Forest for GM data, in order to identify the most relevant species. In Figure 3 we show the Mean Decrease Gini value in a log-normal scale, which represents the relative importance for every identified GM species. In the case of protein, from the total species identified only 740 contribute informationally to the Random Forest, while for the case of lipids it turned out to be a similar number of species, resulting in 721 (Fig. 3).

**Figure 3.**
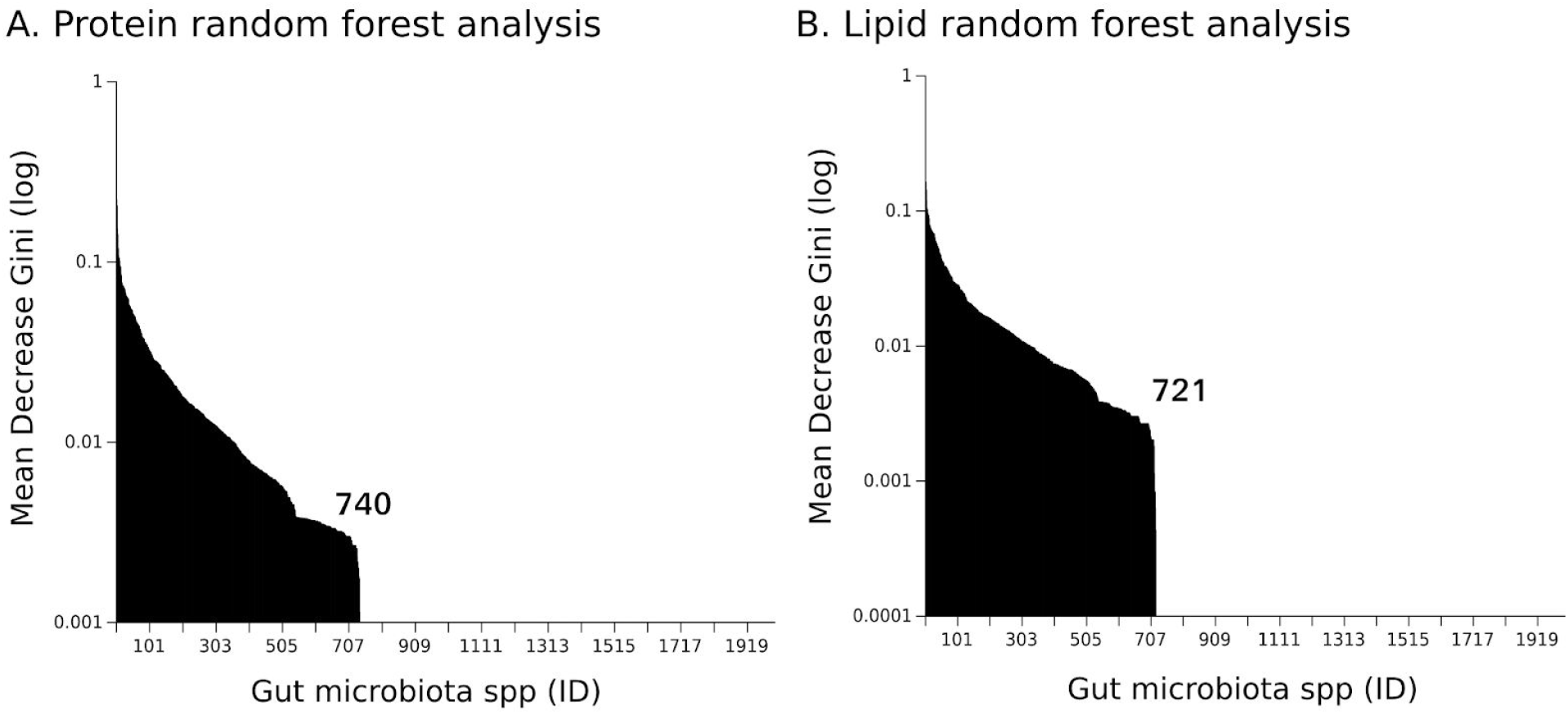
Results from Random Forest of Mean Decrease Gini values that tell us the relative importance of different species to the classification processes into Low Vs High intake for both protein and lipid intake.

Similar to brain connectivity, GM connectivity is also diminished with a low protein and lipid intake. In Figure 4 we show GM connectivity calculated as the total weight of the corresponding Minimum Spanning Tree of Low Vs High intake of both protein and lipids intake.

**Fig 4.**
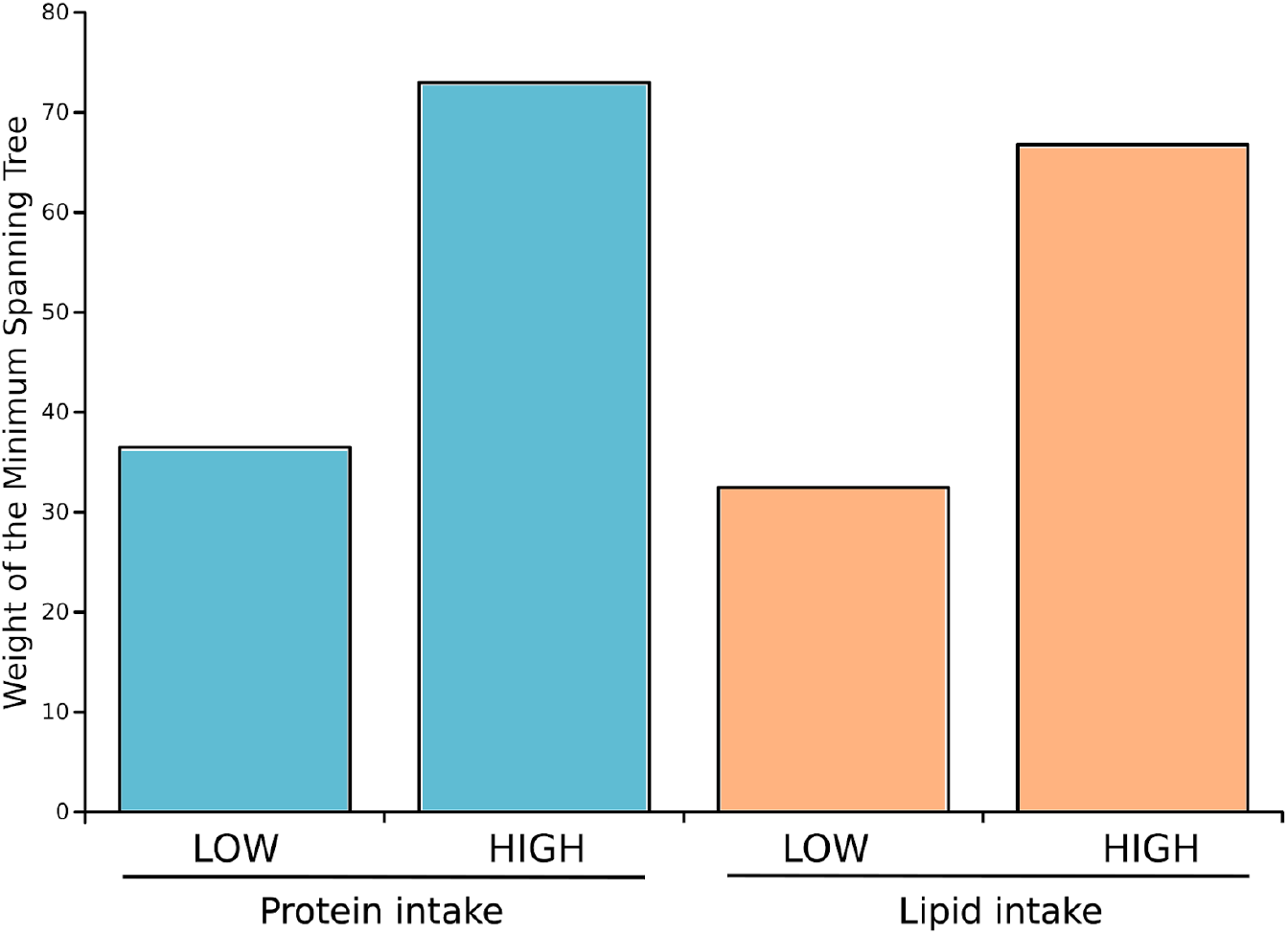
Weight of the MST as a measure of total structural and functional network connectivity (the higher the value, the higher connectivity), for both protein and lipid intake, which is clearly greater in the HIGH intake treatment.

## Discussion

Here we used novel methodologies (Saba et.al. 2019; Ramírez-Carrillo et al. 2020) for analyzing both brain cortex activity and GM as complex systems. Our results show an important influence of animal proteins and lipids consumption over these systems.

Complex systems tend to exhibit scale invariance and a balance between self-organization (order) and emergence (randomness), called criticality. This feature that might be universal in complex systems, implies that systems under this special state, reaches the highest level of computing capacity and they achieve an optimal balance between robustness and flexibility. This so-called Criticality Hypothesis is supported by numerous results from cell biology, physiology to neurosciences (Kiyono, 2001; Goldberger, 2002; Roli et.al., 2018). Of particular interest is the work of Cocchi and co-workers (2017) that summarize evidence about the brain being in criticality, and this as a marker of health.

Even more, we know that important informational quantities maximize under criticality as Complexity, Fisher information or MI (Matsuda et.al. 1996; Pineda et.al. 2019; López-Corona and Padilla, 2019) which conferred the systems with optimum capabilities to construct internal representations which are essential to perceive and respond to environmental changes and to interact with other similar entities. The better these representations (that extract, summarize, and integrate relevant information) the greater competitive advantage either from a cognitive and ecosystemic perspective (Hidalgo et.al. 2014). In this way, as in criticality MI is maximum, a diminish it this quantity implies a departure from criticality and consequently also the loss of this special characteristics, also related with the way systems responds to perturbations; its Antifragility, a property that enhances the system’s functional capacity to respond to external perturbations (Taleb, 2018).

In this manner, the use of MI as presented in this work is not only, as we have said before, better in terms of the mathematical properties of entropy-based metrics as MI; its use is also supported by empirical results; and finally, as we discussed in the previous paragraphs, it makes sense in a Criticality-Antifragility framework. But more important is that this framework has been recently used also in ecology (Ramírez-Carrillo et.al. 2018; Equihua et.al. 2020) and so we can work in a unified manner both microbiota and brain data.

Our results suggest that brain connectivity measured by the MST, or critical backbone of information flow, in the MI network of the resting-state EEG activity, improve for all brain bands under open eyes condition (cognitively demanding) when protein or lipids intake increase (with exception of Beta that remains unchanged for lipid intake) in the indigenous population, which is known to be a dietary deficiency condition in terms of national mean intake values. We observed the same effect for the GM network connectivity which also diminishes under the low intake of protein or lipids. Interestingly enough, from an ecological perspective, a major level of connectivity dissipates the effect of perturbations in the distribution of species and enhances stability, so as suggested by Equihua and co-workers (2020) this would lead to a loss in GM antifragility. Then, under a low nutrient diet GM, the brain or both could get fragilize leading to a systemic loss of health, with potential long term effects. As discussed earlier, a decrease in connectivity measured using MI would mean a relative departure from criticality and consequently a possible loss in cognitive capacities. In this sense, although we did not measure any cognitive process in this indigenous community, the resting-state brain activity we obtained is an important indicator of cerebral self-modulation, a unique feature of the cerebral cortex that ensures that information flows across the entire neural network, ensuring that the information flows to the right place at the right time (Buzsáki, 2007). This organized activity promotes an electrical coupling that establishes temporal windows in which neurons respond to certain stimuli facilitating or limiting the synaptic transmission for different cognitive processes (Engel Fries and Singer, 2001; Fricker and Miles, 2001; Singer, 2011). Hence, an important prospect for this study is to evaluate whether these network connectivity measures, obtained from the resting-state cerebral activity are in fact, associated with demanding cognitive performance, such as working memory or executive functions.

Finally, although the current population is a good experimental model, because of the small number of individuals studied we could not separate the effect of sex (male/female) that could be important. In future work, we want to test differences between age groups and contrast with other populations with a clear difference in lifestyles such as urban population under poverty and not.

## Acknowledgments

This project was funded by UNAM-PAPIIT [grant numbers IA209416, IA207019], CONACYT Ciencia Básica [grant number 241744] and Instituto de Ecología-UNAM [L.I.F].

We are grateful to Margarita Muciño, Julio Naranjo, and Diego Hernández-Muciño from Xuajin Me’Phaa a non-governmental association, for their help in the liaison with the Me’Phaa community, their help in sample collection and logistic. We also are grateful to Mariana Martinez and Santiago Martinez Correa for their contribution in DNA amplification.

